# *Sp8* and *Sp6* regulatory functions in the limb bud ectoderm

**DOI:** 10.1101/2020.02.26.965178

**Authors:** Alejandro Castilla-Ibeas, Sandra Moreu, Rocío Pérez-Gómez, Laura Galán, Víctor Campa, Juan F. López-Giménez, Alvaro Rada-Iglesias, Maria A. Ros

## Abstract

*Sp8* and *Sp6* are two closely related *Sp* genes expressed in the limb ectoderm, where they regulate proximo-distal and dorso-ventral patterning. Genetic studies in mice have shown that they act in a dose-dependent manner, with *Sp8* exerting a substantially stronger effect than *Sp6*. Here, we integrate ChIPmentation-seq and RNA-seq analyses to elucidate their genome-wide regulatory networks and mechanisms of action. Our results show that Sp8 has a dual mode of action either directly binding GC-rich motifs or indirectly binding AT-rich motifs through Dlx. In contrast, Sp6 primarily acts through AT-rich motifs, via interaction with cofactors such as Dlx5. Both factors regulate key components of the Wnt, FGF, and BMP signaling pathways, central to limb patterning, and display both cooperative and independent roles. We further show that Sp8 and Sp6 can form homo- and heterodimers, as well as interact with Dlx5, revealing a complex regulatory network. Our findings provide molecular insight into limb morphogenesis and have potential implications for understanding congenital limb malformations.

## INTRODUCTION

The developing vertebrate limb has long been an excellent system for studying the mechanisms involved in pattern formation and morphogenesis, and more recently in transcriptional regulation. Limb development begins with the emergence of the limb progenitor cells, which originate from the somatopleura of the lateral plate mesoderm through an epithelial-to-mesenchymal transition. Subsequent limb outgrowth and patterning rely on the interactions between these progenitor cells and the overlying ectoderm. One of the earliest interactions induces *Fgf8* expression in the ectoderm, leading to the formation of the apical ectodermal ridge (AER), a crucial signaling center that supports the survival and proliferation of the limb progenitor cells (Fernandez-Teran & Ros, 2008; Tickle, 2015). Simultaneously, *En1* is activated in the ventral ectoderm, restricting *Wnt7a* expression to the dorsal ectoderm and thereby establishing distinct dorsal-ventral (DV) gene expression domains essential for proper DV limb patterning. In the dorsal ectoderm, *Wnt7a* promotes limb bud dorsalization by inducing the expression of the homeobox gene *Lmx1b* in the underlying mesoderm (Fernandez-Teran & Ros, 2008; Tickle, 2015).

Elegant studies demonstrated that the activation of *Fgf8* and *En1* in the limb ectoderm depends on active Wnt/βcatenin and Bmp signaling (Barrow et al., 2003; Kyung Ahn et al., 2001; Soshnikova et al., 2003). The transcription factors (TF) Sp6 and Sp8 act downstream of these pathways as critical mediators of *Fgf8* and *En1* induction (Haro et al., 2014) but the underlying molecular mechanisms remain incompletely understood. Understanding the mechanisms that lead to the induction and maintenance of the AER as well as DV polarity is of maximum interest not only for advancing our understanding of limb development but also for informing strategies in tissue regeneration and repair.

Sp6 and Sp8 belong to the Specificity protein/Krüppel-like (Sp/Klf) family found across almost all metazoan species (Presnell et al., 2015; Schaeper et al., 2010; Suske, 1999). The Sp family is characterized by a highly conserved carboxyterminal DNA binding domain consisting of three consecutive C2H2-type zinc finger (ZF) motifs and a more variable N-terminal region (Suske et al., 2005). The buttonhead box and the Sp box are also conserved structural domains characteristic of these proteins.

During limb development, Sp6 and Sp8 are exclusively expressed in the ectoderm where their cooperative function is necessary for proximo-distal (PD) and DV patterning (Bell et al., 2003; Haro et al., 2014; Talamillo et al., 2010; Treichel et al., 2003). The analysis of *Sp8;Sp6* double mutants revealed that they act in a dose-dependent manner, with their combined absence leading to tetra-amelia (total loss of limb formation) due to a failure in *Fgf8* activation. Partial loss of *Sp6* and *Sp8*, such as in mutants with a single functional *Sp8* allele (*Sp6⁻/⁻*;*Sp8⁺/⁻*), leads to split-hand/foot malformation (SHFM), characterized by irregular and immature AER formation and double dorsal digit tips (Haro et al., 2014). A reasonable conclusion of these studies, supported by some *in vitro* assays (Sahara et al., 2007), was that Sp8 and Sp6 were transcriptional activators of *Fgf8* and *En1* linking PD and DV patterning. However, the full repertoire of Sp6 and Sp8 target genes, whether shared or distinct, as well as the molecular mechanisms and genomic context orchestrating their regulation are not completely known.

Here, we combined genome-wide chromatin occupancy (ChIPmentation-seq) with transcriptomic profiling (RNA-seq) to identify shared and specific Sp8 and Sp6 direct targets in the limb ectoderm in vivo. We found that Sp8 and Sp6 act mainly as transcriptional activators via distal enhancers with many co-occupied regulatory regions, supporting a cooperative function. However, within these shared regions, Sp8 appears to be functionally more dominant. In addition, Sp8 and Sp6 also have distinct individual functions; for example, Sp8 uniquely regulates *Fgf8*, while Sp6 plays a specific role in epithelial organization.

An additional key finding of this study is the mechanistic divergence between Sp8 and Sp6 modes of action. While Sp8 directly binds the canonical GC-box motif, a hallmark of Sp transcription factors, Sp6 shows little evidence of direct DNA binding and instead primarily targets AT-rich motifs via cofactors such as Dlx5. Sp8 can also bind AT-rich motifs indirectly, showing a dual DNA-binding mechanism. The co-expression of several *Dlx* genes with *Sp8* and *Sp6* in the developing limb ectoderm, along with evidence from co-IP and EMSA assays, supports a potential functional interaction between Sp and Dlx factors. We propose that the Sp/Dlx interaction should be considered when interpreting the phenotypes of Sp and Dlx mutants and their association with congenital malformations.

## RESULTS

### Generation of a *Sp8*-tagged Knock-in Mouse

The genome-wide binding of Sp8 has not been previously mapped in the limb ectoderm or in any other cellular context. This is probably due to the lack of commercially available ChIP-grade antibodies for Sp8. To overcome this limitation and, thus, globally identify Sp8 binding sites in the limb ectoderm, we generated a knock-in (KI) mouse in which the endogenous *Sp8* gene was tagged with three copies of the FLAG (FL) epitope using homologous recombination (*Sp8FL*; Fig. 1A). This epitope was incorporated in frame at the C terminus of Sp8 protein, a strategy that has been widely used in ChIP-seq studies (Osterwalder et al., 2014; Vokes et al., 2008). Mice homozygous for the *Sp8FL* allele were viable and fertile and displayed no obvious phenotype indicating that the tagged protein was fully functional (Fig. 1B). Accordingly, the detection of the Sp8FL protein in immunofluorescence assays using the αFLAG M2 antibody (Ref F1804, Sigma) was similar to that of the endogenous protein detected with an αSp8 antibody (Ref 104661, Santa Cruz) (Fig. S1A). The αFLAG antibody also readily detected the Sp8FL protein in Western blots (Fig. S1B). Thus, the *Sp8FL* KI model provides a useful resource for future studies requiring detection of Sp8.

**Figure 1.**
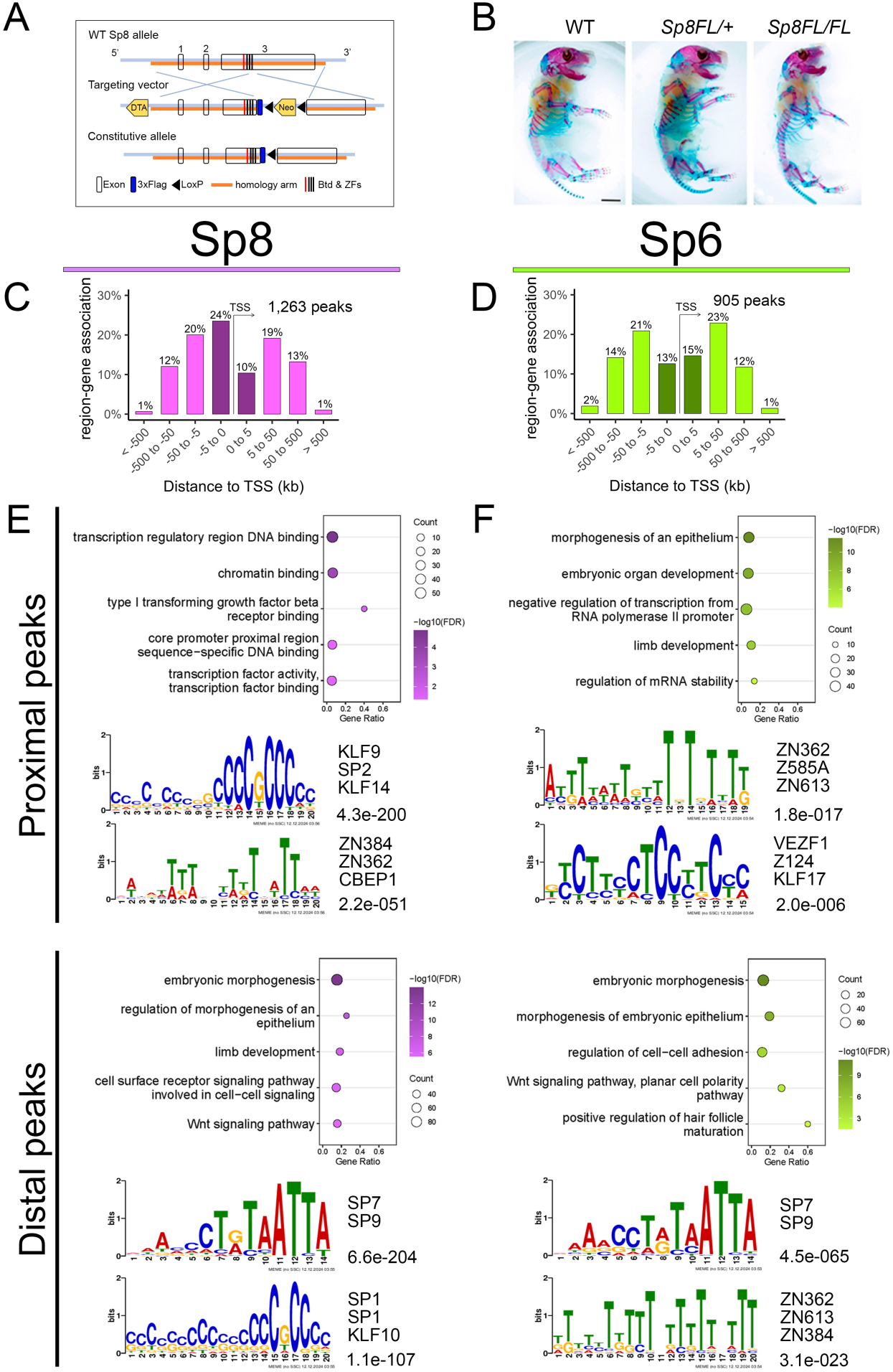
Genome-wide analysis of Sp8 and Sp6 binding profiles and functional peak distribution. **(A)** Scheme showing the generation of the *Sp8-3xFLAG* (*Sp8FL*) allele by homologous recombination (homology arms in orange). Three copies of FLAG (dark blue) were introduced at the 3’-terminal end of the *Sp8* gene before the STOP codon TGA (middle of third exon). Numbers 1, 2, 3 indicate *Sp8* exons (white). **(B)** Skeletal preparations of WT, and Sp8FL hetero and homozygote newborn mice. **(C-D)** Distribution of Sp8 (C) and Sp6 (D) peaks relative to their distance to the nearest TSS. **(E-F)** Analysis of enrichment of GO:biological process (top) and de-novo discovered motifs (bottom) for Sp8 **(E)** and Sp6 **(F)** separately in proximal (<5kb) and distal peaks (>5kb). Top 2 motifs with best matches of known motifs and e-values are shown.

### Global Mapping of Sp8 and Sp6 Binding Sites in the Limb Ectoderm

To investigate the genome-wide occupancy of Sp8 and Sp6 in limb ectodermal cells, we used ChIPmentation-seq. This method, which combines chromatin immunoprecipitation (ChIP) with tagmentation, has proven highly efficient with low numbers of starting cells (Schmidl et al., 2015), enabling us to reduce the initial sample size to 50 ectodermal hulls of E10.5 forelimb buds (approximately 7.5×10^5^ cells). The ectodermal sheets were isolated using mild trypsin digestion (see M&M). Limb buds from embryonic day (E) 10.5 were selected, as this stage corresponds to a fully matured AER and coincides with the onset of both *Sp8* and *Sp6* null phenotypes (Bell et al., 2003; Fernandez-Teran & Ros, 2008; Talamillo et al., 2010; Treichel et al., 2003). The genome-wide binding of Sp8 was mapped using homozygous *Sp8FL* embryos and the αFLAG. For Sp6, wild-type (WT) embryos and the αSp6 antibody (Ref 21234-1-AP, Proteintech) were used. We previously validated the αSp6 antibody through immunofluorescence (Fig. S1C) and confirmed its specificity in Western blots by showing the absence of a band in *Sp6*-null limb buds and no off-target bands from other Sp proteins (Fig. S1D).

Following sequencing, analysis of the ChIPmentation datasets resulted in the identification of 1,263 high-confidence Sp8 binding sites (Source Data 1) and 905 high-confidence Sp6 binding sites (Source Data 1). The quality and specificity of the identified Sp8 and Sp6 peaks was supported by (i) their high evolutionary conservation (Fig. S1E), (ii) reproducibility in an independent biological ChIPmentation replicate (Fig. S1F), (iii) high correlation between the two replicates (Fig. S1G), and (iv) overlap with highly accessible regions in the limb bud ectoderm identified by single cell ATAC-seq (Desanlis et al., 2020) (Fig. S1H).

The *de novo in-silico* analysis of sequence motifs in the Sp8 peaks using the MEME Suite (Bailey et al., 2015) revealed significant enrichment of a CG-rich/Sp motif (E-value: 3.3e-161) and an AT-rich/Sp7 motif (E-value: 1.2e-154) as the most over-represented motifs. Another C-rich motif associated with ZN transcription factors was also modestly enriched (Fig. S1I). The presence of the Sp consensus binding sequence, the GC box: 5’-CCGCCC-3’, was expected as it represents the conserved DNA recognition motif shared by Sp family members (Suske, 1999). The AT-rich/SP7 motif has previously been shown to enable Sp7 to bind DNA indirectly via interaction with Dlx5 (Hojo et al., 2016), suggesting that Sp8 may also interact with Dlx. Interestingly, Sp9, another Sp family member, has also been shown to exhibit enrichment of both AT-rich motifs (Xu et al., 2018) and GC boxes (Catta-Preta et al., 2025).

In contrast, the *de novo in-silico* analysis of Sp6 peak sequences did not identify the canonical GC-box Sp family binding motif. Instead, there was significant enrichment of the AT-rich/Sp7 motif (E-value 3.9e-066), along with other motifs associated -with zinc finger transcription factors (Fig. S1I). The absence of the Sp consensus motif is consistent with findings from a previous ChIP-seq study in developing teeth (Rhodes et al., 2021), which identified a core motif (CTg/aTAATTA) closely resembling the Sp7 binding motif in bone (Hojo et al., 2016) and the motif found in the present study. These results suggest that Sp6 primarily regulates gene expression through indirect mechanisms, involving Dlx and other co-factors, at least in its major sites of expression, such as the tooth and limb.

### Distinct Functional and Genomic Features of Proximal and Distal Sp6 and Sp8 Regulatory Elements

Being confident about the quality of our Sp8 and Sp6 binding profiles, we then mapped the locations of the peaks with respect to the nearest transcription start site (TSS). While both transcription factors bind proximal (<5kb from TSS) and distal (>5kb from TSS) cis-regulatory element contexts, there is a moderate preference for distal regions, 66% for Sp8 (Fig. 1C) and 73% for Sp6 (Fig. 1D) and, therefore, long-range regulatory interactions.

To explore potential functional differences based on whether binding occurred in proximal or distal regulatory regions, we performed separate *in silico* analyses of proximal versus distal Sp8 and Sp6 peaks. Functional annotation of Sp8 peaks using GREAT (McLean et al., 2010) revealed that proximal binding was primarily associated with general biologic processes, such as “transcription regulatory region DNA binding” and “transcription factor activity” (Fig. 1E, Source Data 2). Conversely, distal binding predominantly associated with processes related to “embryonic morphogenesis”, “limb development” and “Wnt signaling pathway”. Moreover, we also examined whether Sp8 recognized different DNA sequences when binding proximal or distal regulatory sites. *de novo* motif analysis revealed that both proximal (E-value 4.3e-200) and distal (E-value 1.1e-107) Sp8-bound elements were enriched in the Sp family consensus GC-rich motif (Fig. 1E). In addition, the AT-rich/Sp7 motif was also enriched in the distal set of peaks where it was the most prominent (E-value 6.6e-204). Thus, Sp8 may use distinct regulatory mechanisms when operating at either promoters or enhancers.

A similar analysis of Sp6 peaks revealed enrichment in terms related to embryonic and epithelial morphogenesis in both proximal and distal elements (Fig. 1F). Consistent with the global peak analysis, the consensus Sp motif was absent from both proximal and distal peaks, whereas the Dlx core motif was the most enriched in distal peaks (E-value 4.5e-065). The smaller number of peaks in the proximal set limited the robustness of the analysis for this group (Fig. 1F, Source Data 1).

Together, our results suggest that Sp8 operates through two distinct functional modes. First, directly binding DNA at Sp consensus sequences, predominantly located at proximal promoters, to regulate general biological processes. Second, indirectly interacting with DNA via binding partners such as Dlx5, and potentially other Dlx family members, at distal putative enhancers, thereby controlling genes involved in specific limb developmental functions. In contrast, Sp6 appears to function primarily through indirect mechanisms involving Dlx and other cofactors, independent of genomic location.

### Shared and Distinct Binding Dynamics of Sp8 and Sp6 During Limb Development

Having mapped the individual binding profiles of Sp8 and Sp6 across the genome and considering their dose-dependent cooperation during limb development (Haro et al., 2014), we analyzed the extent of overlap in their binding sites. Surprisingly, we found minimal direct overlap among the Sp8 and Sp6 peaks identified by MACS2, with only 185 peaks being shared between the two factors. This limited overlap could be attributed to differences in ChIP-seq sensitivity, potentially obscuring the detection of combinatorial binding events (Catta-Preta et al., 2025). Indeed, ChIPmentation tracks revealed substantial signal overlap between Sp8 and Sp6, suggesting a more extensive co-occupancy. Thus, to further characterize Sp8/Sp6 cooperative binding, we performed unbiased k-means clustering of ChIPmentation signals with the merged set of peaks which identified three distinct peak categories (Fig. 2A; Source Data 5): (i) peaks with high signal overlap between Sp8 and Sp6 (607 peaks, 31.58%), (ii) peaks predominantly bound by Sp8 (803 peaks, 41.77%), and (iii) peaks predominantly bound by Sp6 (512 peaks, 26.33%).

**Figure 2.**
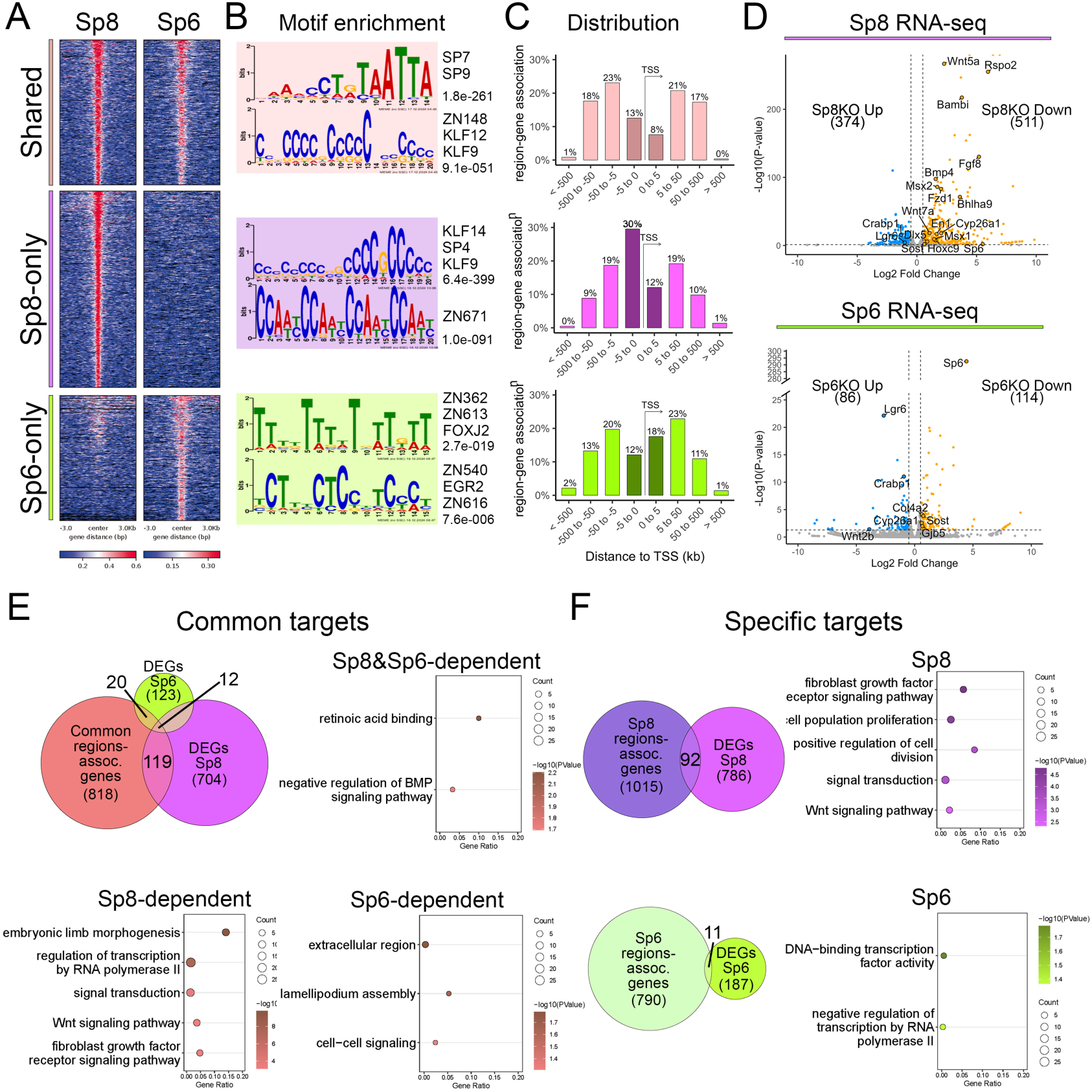
Genome-wide analysis of shared and specific Sp8 and Sp6 binding sites. **(A)** Heatmaps of the three peak clusters, bound by both Sp8 and Sp6 (Shared), by Sp8 only, and by Sp6 only, obtained through k-means clustering of the merged peak set. **(B)** Top 2 most enriched motifs obtained by MEME-ChIP *de novo* motif discovery tool for each cluster. Best matches of known motifs and e-values on the right side. **(C)** Distribution of peaks relative to their distance to the nearest TSS for each cluster. **(D)** Volcano plots of the differential expression analysis obtained from the comparison between WT and *Sp8* homozygous mutants (top) and between WT and *Sp6* homozygous mutants (bottom) with the number of DEGs and selected genes indicated. **(E)** Analysis of commonly bound regions. Venn plot of the intersection of the region-associated genes with Sp8 and Sp6 DEGs and plots of GO:BP terms enriched in each corresponding subcategory as indicated. **(F)** Analysis of specifically bound regions. Venn plot of the intersection of the corresponding region-associated genes with Sp8 DEGs and Sp6 DEGs and plots of GO:BP terms enriched in the corresponding category.

MEME-ChIP analysis (Fig. 2B) of regions co-occupied by Sp8 and Sp6 revealed strong enrichment for the AT-rich motif (E-value 1.8e-261), with a much less significant enrichment for the GC-box motif (E-value 9.1e-051). In contrast, regions bound exclusively by Sp8 showed a strong preference for the GC-box motif (E-value: 6.4e-399), while regions bound exclusively by Sp6 were enriched for ZN motifs, albeit with lower statistical significance. These results suggest that Sp8 and Sp6 employ distinct binding strategies depending on whether they act together or independently. Specifically, their co-occupancy correlates with enrichment in AT-rich motifs, while Sp8 binding alone correlates with GC-rich motifs, and Sp6 binding alone appears to be linked to partnerships with ZN transcription factors. The presence of GC-box and AT-rich motifs was also analyzed individually for every peak (Fig S2A) with FIMO algorithm (Grant et al., 2011). Consistent with previous results, a significant proportion of common peaks (39%) contained only AT-rich motifs, suggesting that there are genes that are regulated by Sp8/6 solely through AT-rich motifs.

Finally, we analyzed the genomic distribution of the three peak categories (Fig. 2C). Co-occupied peaks and those exclusive to Sp6 were strongly biased toward distal elements (79% and 70%, respectively). By contrast, Sp8-specific peaks exhibited a more even distribution, with 58% in distal regions, suggesting that, in the absence of Sp6, Sp8 has relatively greater engagement at promoter-proximal sites.

### Impact of Sp8 and Sp6 Loss on the Transcriptome of Limb Bud Ectodermal Cells

To gain further insights into the developmental program regulated by Sp8 and Sp6 in the limb ectoderm, we compared the expression profiles of the E10.5 WT and *Sp8* or *Sp6* deficient limb bud ectoderms. For this, we used a robust RNA-seq protocol for low-abundance RNA (see M&M for details).

Differential gene expression analysis identified 885 differentially expressed genes (DEGs) in *Sp8*-null embryos (Fig. 2D; Source Data 3), with 511 downregulated and 374 upregulated, suggesting that Sp8 acts as both an activator and a repressor, with a slightly stronger role as an activator. Notably, all genes previously reported to be affected by Sp8 loss, including *Fgf8, En1, Msx2, Rspo2,* and *Sp6* (Bell et al., 2003; Haro et al., 2014; Treichel et al., 2003), were among the DEGs, validating the quality and accuracy of our RNA-seq data.

A similar analysis in Sp6-deficient limb ectoderms identified 200 DEGs, with 114 downregulated and 86 upregulated (Fig. 2D; Source Data 3). This suggests Sp6 acts as both an activator and a repressor, with a slightly stronger role as an activator. Consistent with the limb *Sp6*-null phenotype (Talamillo et al., 2010), genes altered in the absence of *Sp6* included key modulators of the Wnt/β-catenin signaling pathway, such as *Sost*, as well as extracellular matrix components such as *Col4a2*, a major constituent of the basement membrane.

Gene Ontology (GO) enrichment analysis of the DEGs using DAVID revealed distinct biological processes influenced by these transcription factors. Sp8 DEGs are enriched in “multicellular organism development,” “embryonic limb morphogenesis,” “BMP signaling pathway,” and “fibroblast growth factor receptor signaling pathway”, indicating an important function during limb morphogenesis. Sp6 DEGs are linked to “extracellular matrix organization,” “canonical Wnt signaling pathway,” and “cell adhesion,” (Fig. S2B, Source data 3). Thus, while both Sp8 and Sp6 influence key biological processes in the limb ectoderm, the loss of Sp8 has a greater transcriptomic impact than the loss of Sp6, consistent with their mutant phenotypes.

### Identification of Direct Sp8 and Sp6 Targets in the Limb Bud Ectoderm

To distinguish direct from indirect Sp8 and Sp6 target genes and uncover the transcriptional regulatory networks they govern, we intersected the list of genes assigned to each binding region category (co-occupied, primarily bound by Sp8 and primarily bound by Sp6) with the list of DEGs identified in *Sp8* and *Sp6* null mutants (Fig. 2E; Source Data 5).

Among the genomic regions co-occupied by Sp8 and Sp6, we identified 24 peaks linked to 12 DEGs in both *Sp8KO* and *Sp6KO* mutants (Fig. 2E and Source Data 5). This set included *Cyp26a1* and *Crabp1*, regulators of retinoic acid metabolism, which showed complex regulation: *Cyp26a1* was downregulated in *Sp8KO* but upregulated in *Sp6KO*, while *Crabp1* was downregulated in both mutants, suggesting a complex modulation of RA signaling in the limb ectoderm (Niederreither & Dollé, 2008). It also included *Sost*, a negative modulator of Wnt signaling (Collette et al., 2010; Semënov et al., 2005) expressed in the limb bud ectoderm but not the AER (Fernandez-Guerrero et al., 2021) also implicated in BMP regulation (Winkler et al., 2003). GO analysis using DAVID confirmed enrichment for terms such as “retinoic acid binding” and “negative regulation of BMP signaling pathway” (Fig. 2E).

A subset of co-occupied regions was associated with genes differentially expressed in only one mutant, despite potential co-regulation, indicating differential dependence on Sp8 or Sp6 (Fig. 2E). Among genes co-regulated but specifically dependent on Sp8, we identified 119 genes (linked to 140 peaks), including key limb development genes (*Bambi*, *En1*, *Wnt7a*, *Hoxc9*, *Rspo2*, *Fzd1*, *Msx1, Msx2* and *Sp6*), all positively regulated by Sp8. GO analysis highlighted terms related to “embryonic limb morphogenesis” and “Wnt signaling pathway”. Among genes specifically dependent on Sp6, we found 20 genes (22 peaks), including typical ectodermal genes (*Col4a2* and *Gjb5*). Corresponding GO terms included “cell-cell signaling” and “extracellular region”. A substantial number of putative shared targets showed no expression changes in single mutants, suggesting mutual compensation or nonessential roles. Transcriptomic analysis of *Sp8/Sp6* double mutant limb ectoderms could clarify between these scenarios; however, their severe, early-onset limb defects limited the collection of ectoderm suitable for high-quality profiling.

Sp8 specific peaks were linked to 92 DEGs in *Sp8*-null limb buds representing regions associated with Sp8 targets not regulated by Sp6. These included key genes involved in limb development, most notably *Fgf8*, as well as *Kremen1* and *Prickle1*, all of which positively regulated by Sp8. GO analysis revealed enrichment for “fibroblast growth factor receptor signaling pathway,” “positive regulation of cell division,” and “positive regulation of ERK1 and ERK2 cascade” consistent with Sp8’s role in regulating AER function (Fig. 2E). Conversely, Sp6 specific peaks were associated with 11 DEGs in *Sp6*-null mutants, none of which with known, relevant functions in limb development. The associated GO terms were less specific, including “negative regulation of transcription” underscoring Sp6’s more limited regulatory specificity when acting independently of Sp8.

Finally, we note that some targets are associated with multiple regulatory elements across categories (Fig. S2D). 25 genes are regulated by shared and by Sp8-specific regions. For example, *Wnt5a* is linked to two distal regulatory elements co-bound by Sp8 and Sp6 and a third distal region bound exclusively by Sp8, suggesting complex regulation (Fig S2E).

### Validation of Sp8 and Sp6 Target Genes

To confirm direct Sp8 and Sp6 regulation and assess spatial expression patterns, we performed in situ hybridization on direct targets selected for their relevance to limb development. In addition, transgenic reporter assays were performed in selected cases to assess enhancer activity.

*Sost* was selected from the set of Sp8 and Sp6 common target genes that require the input of both TF for normal expression (Fig 3A). Sp8 and Sp6 bind to an ATAC-seq accessible region within the first intron of *Sost*. In situ hybridization on tissue sections revealed that *Sost* was expressed in the WT non-AER ectoderm and markedly downregulated in *Sp8* and *Sp6* mutants, more severely in *Sp8*-null mutants, and completely lost in double mutants. These results confirm *Sost* as bona fide direct target under cooperative regulation by Sp8 and Sp6.

**Figure 3.**
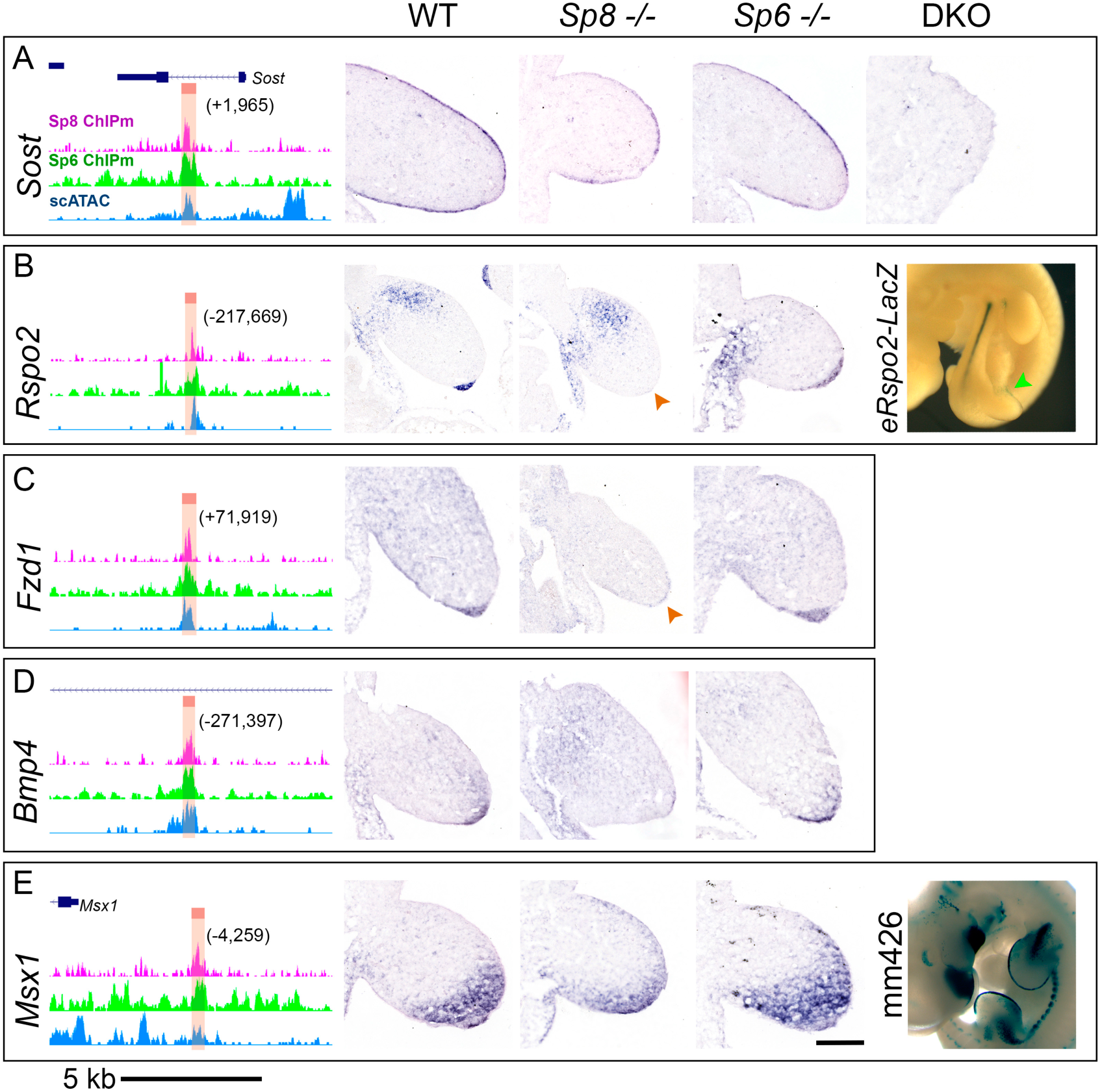
In situ hybridization validation of Sp8 and Sp6 target genes. (**A-E**) For each of the indicated genes, a screenshot of the UCSC genome browser displaying Sp8 (pink) and Sp6 (green) ChIPmentation tracks and the scATAC-seq accessibility in the ectoderm (blue) is shown. The Sp8/Sp6-associated regulatory modules are highlighted in orange. On the right ISH images of E10.5 longitudinal limb bud sections of the genotypes indicated at the top. LacZ reporter activity is shown for the *Rspo2* (B, right; *eRspo2-LacZ*) and the *Msx1* (E, right; mm426 from VISTA), putative enhancers. Genomic distances to each gene’s transcription start site are indicated in parentheses. Scale bar in genomic tracks = 5 kb. ISH scale bar=100 µm

Among the extensive list of targets co-regulated by Sp8 and Sp6 but dependent solely on Sp8 for normal expression, we selected *Rspo2*, *Fzd1*, *Bmp4*, *Msx1*, *En1*, *Wnt7a*, and *Bhlha9*, all critical for limb development, for detailed analysis. ISH confirmed specific loss of *Rspo2* in the AER and *Fzd1* expression in the ectoderm of *Sp8-*null mutant with no obvious changes in *Sp6* mutants (Fig 3B-C, orange arrowheads). However, the putative *Rspo2* enhancer located 217 kb upstream of the TSS, displayed only weak activity in the limb ectoderm (6/15 embryos) in transgenic reporter assays (Fig 3B, green arrowhead). The expression of *Bmp4* and its bona fide targets *Msx1* and *Msx2* were also specifically downregulated in the ectoderm of *Sp8* mutant limb buds but not in *Sp6* mutants (Fig 3D-E and S3B). The *Msx1* assigned peak has already been tested in the VISTA enhancer dataset (Visel et al., 2007) and shows strong activity in the AER. Thus, our results suggest that Sp8 and Sp6 modulate Wnt and Bmp signaling pathways by enhancing Wnt signal reception and Bmp ligands and targets.

Our results also identify *En1* and *Wnt7a*, key regulators of DV limb patterning, as direct targets of Sp8 and Sp6 (Fig 4A-B). *En1* expression, previously shown to be downregulated in *Sp8* and double *Sp8;Sp6* mutants, is controlled by a distal downstream enhancer (Fig. 4A). Transgenic reporter assays confirmed this enhancer’s activity in the AER with high specificity and reproducibility (3 out of 4; Fig. 4A). In *Sp8* mutants and *Sp8;Sp6* double mutants, *Wnt7a* expression has been reported to expand into the ventral ectoderm as a consequence of *En1* loss (Haro et al., 2014). Here we show that *Wnt7a* is also directly and positively regulated by Sp8/Sp6, potentially through a distal downstream enhancer. Accordingly, in addition to the expansion of its expression domain, the overall expression level of *Wnt7a* is reduced in the ectoderm of *Sp8* mutants (Fig. 4B).

**Figure 4.**
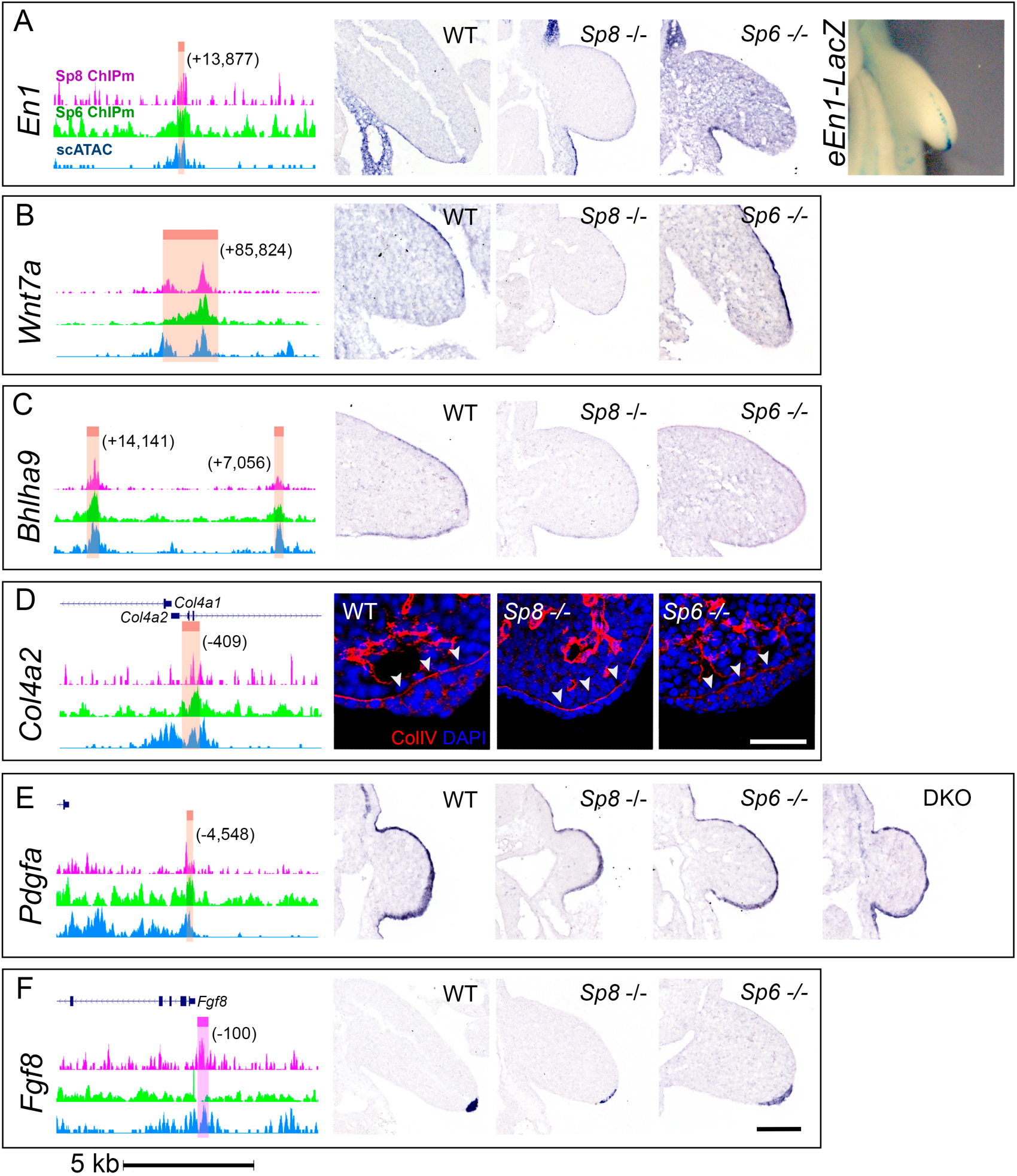
In situ hybridization validation of Sp8 and Sp6 target genes (Continuation). (A-F) Panels as in Fig. 3. LacZ reporter activity is shown in **(A)** for the *En1* enhancer. IF for Col4a2 showing significant reduction in the basement membrane (indicated by white arrowheads) of *Sp6* mutants is shown in **(D)**. Scale bar in genomic tracks = 5 kb. IF scale bar=50 µm. ISH scale bar=100 µm

*Bhlha9* (*Fingerin*), is a basic helix-loop-helix transcription factor involved in limb development that in the mouse has a phenotype similar to that of Sp6. In humans, *BHLHA9* is linked to split-hand/foot malformation with long bone deficiency (SHFLD3) a phenotype resembling *Sp6;Sp8* compound mutants retaining a functional copy of *Sp8*. The ISH analysis (Fig 4C) confirmed a marked downregulation in Sp8 mutants.

Among the common targets dependent solely on Sp6 for normal expression, we selected *Col4a2*. Immunofluorescence analysis for Collagen type IV confirmed a marked reduction in the basement membrane in the absence of *Sp6* (Fig. 4D).

Among the shared target genes that do not exhibit changes in any of the mutants, we examined *Pdgfa* (Fernandez-Guerrero et al., 2021). No alteration in expression was observed (Fig 4E), suggesting that Sp8 and Sp6 binding may not play a regulatory role in this context.

Among the genes exclusively regulated by Sp8 we selected *Fgf8* because of its importance in limb development. The Sp8-peak assigned to *Fgf8* was in the gene promoter (-326 bp) and contained several Sp1-like binding sites supporting direct regulation by Sp8. Curiously, no Sp6 binding was detected in this region. In *Sp8* mutant limb buds, *Fgf8* is initially activated but its expression decays concomitantly with the onset of the phenotype indicating that this regulatory region is probably involved in maintenance of expression (Fig. 4F) (Bell et al., 2003; Treichel et al., 2003). In contrast *Fgf8* expression in *Sp6* mutants is not altered in its level of expression but in the domain of expression consistent with the mild double ridge phenotype typically observed in these mutants.

Other genes important for limb development, included *Wnt5a*, *Bambi*, *Dlx1, Dlx2*, *Dlx4* and *Sp6* some of them also validated by ISH (Fig. S3B-D; Source Data 5). Collectively, these findings demonstrate that Sp8 and Sp6 directly regulate essential genes in the limb bud ectoderm, highlighting the cooperative as well as independent roles of these transcription factors.

### Distinct DNA-Binding Modes of Sp6 and Sp8

Our data show that Sp6 primarily binds to an AT-rich Dlx motif, rather than the canonical GC-box. This AT-rich motif is also enriched in Sp8-bound regions, suggesting that both Sp6 and Sp8 may operate through a mechanism similar to that used by Sp7 in osteoblasts.

To evaluate the binding affinities of Sp8 and Sp6 for AT-rich and GC-box motifs, we performed an electrophoretic mobility shift assay (EMSA) using two constructs: one containing the AT-rich/Dlx5 motif (Fig. 5A) (Hojo et al., 2016) and another containing the GC-box motif (Fig. 5B) (Vaqué et al., 2005). Neither Sp8 nor Sp6 bound directly to the AT-rich motif, suggesting that their association with this motif is likely indirect and dependent on cofactors (Fig. 5A). In contrast, Sp8 (FLAG-tagged) exhibited strong and specific binding to the GC-box, while Sp6 binding was barely detectable, similar to previous findings for Sp7 (Fig. 5B). The Sp8-GC-box interaction was further validated by the supershift in presence of the anti-FLAG antibody (Fig. 5B). These results confirm the distinct modes of action for these transcription factors: whereas Sp8 functions through both direct DNA binding via the GC-box and indirect interactions with other factors, Sp6 predominantly relies on cofactors. Protein expression levels of Sp6 and Sp8 were quantified by Western blot analysis (Fig. S4A).

**Figure 5.**
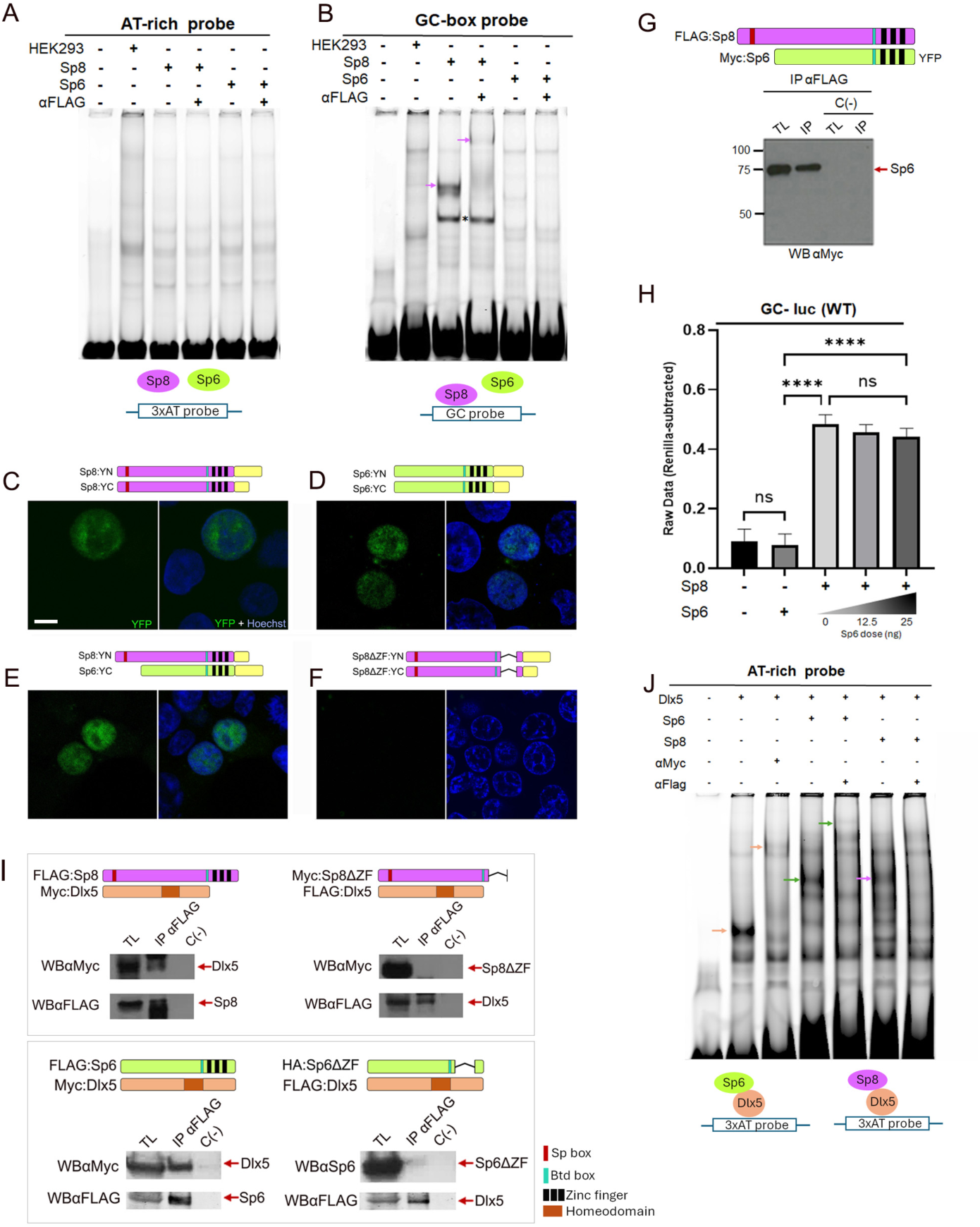
Characterization and functional analysis of Sp8, Sp6 and Dlx5 interactions. **(A)** EMSA using an AT-rich probe shows no specific DNA binding by Sp8 or Sp6. **(B)** EMSA with a GC-box probe shows specific binding of Sp8, indicated by a shift and a supershift upon addition of the anti-FLAG antibody (pink arrowheads, lanes 3–4) but not of Sp6. **(C-F**) BiFC assays shows Sp8 and Sp6 homo and heterodimerization; constructs used shown on top. Note the interactions require the ZF domain (F). Cells were stained with Hoechst to label nuclei. For each experiment, YFP (left) and merged YFP/Hoechst (right) channels are shown. Scale bar: 10 μm. **(G**) Co-IP confirming Sp6-Sp8 heterodimerization. TL: total lysate; C−: negative control; IP: immunoprecipitation. **(H)** Luciferase reporter assay using a GC-box promoter shows that Sp8 activates transcription, while Sp6 does not. Note Sp6 does not interfere with Sp8-mediated activation. Error bars represent standard deviations calculated from biological triplicates. *P < 0.05; ****P < 0.0005. **(I)** Co-IP assays with the constructs indicated on top of each panel show that both Sp8 and Sp6 interact with Dlx5 in HEK293 cells (left panel), and that this interaction requires the ZF domain (right panel). (**J)** EMSA with AT-rich probe shows that Myc-Dlx5 binds the probe (orange arrowhead, lane 2), with supershift upon anti-Myc addition (lane 3). Co-expression with FLAG-Sp6 or FLAG-Sp8 results in additional complexes (green and pink arrowheads, lanes 4 and 6, respectively), which are either supershifted or no longer visible upon addition of anti-FLAG antibody (lanes 5 and 7, respectively), confirming the Dlx-Sp6/Sp8 complex. The legend for the structural domains is shown in panel (I).

### Sp8 and Sp6 form Heterodimers without Affecting Sp8’s Direct DNA Binding

Our findings show that Sp8 and Sp6 co-occupy certain regulatory regions while binding independently to others. Notably, regions bound by both Sp8 and Sp6 are highly enriched in AT-rich/DLX5 motifs whereas regions exclusively bound by Sp8 are primarily linked to GC-rich motifs. This suggests that Sp8 and Sp6 may have distinct DNA-binding preferences, whether functioning independently or together. Considering that Sp transcription factors are known to hetero dimerize, we asked whether Sp8 and Sp6 might physically interact forming heterodimers that could potentially influence their DNA-binding specificity and regulatory activity. To investigate this, we performed Bimolecular Fluorescent Complementation (BiFC) assays to assess direct protein-protein interactions. Sp8 and Sp6 were fused in-frame at their C-terminus to Yellow Fluorescent Protein (YFP) or to the YFP N-terminal (YN; residues 1-172) or C-terminal (YC; residues 173-240) moieties (see Materials & Methods). As expected, both Sp8 and Sp6 are localized to the nucleus (Fig. S4B). BiFC assays revealed that both factors can form homo-and heterodimers in the nucleus (Fig. 5C-E), with this interaction requiring the zinc finger (ZF) domain (Fig. 5F). To further validate Sp8-Sp6 heterodimerization, we performed co-immunoprecipitation (CoIP) assays, which confirmed their direct interaction (Fig. 5G).

Having demonstrated that Sp8 and Sp6 can heterodimerize, we next investigated whether this interaction influences Sp8’s ability to directly bind DNA. Using a luciferase reporter assay with a construct containing four tandem SP1 sites (GC-box) (Vaqué et al., 2005), we found that Sp8 alone significantly activated the luciferase reporter, whereas Sp6 did not (Fig. 5H), consistent with the EMSA results. Co-expression of Sp6 with Sp8 did not alter Sp8’s transcriptional activity, suggesting that (putative) Sp8/Sp6 heterodimers retain Sp8’s ability to directly bind the GC-box, at least under these experimental conditions (Fig. 5H). To confirm specificity, we tested a reporter with mutated GC-box motifs (see M&M) and observed no transcriptional activation, confirming that the effect is GC-box–dependent (Fig. S4C).

### Sp8 and Sp6 as Cofactors of the Dlx Family in Limb Ectoderm Development

Despite the enrichment of the AT-rich/Dlx5 motif in Sp8-and Sp6-bound cis-regulatory regions (Fig. 2C), EMSA assays demonstrated that neither factor directly binds an AT-rich probe (Fig. 5A). This suggests that Sp8 and Sp6 may be recruited to some of their target sites through interactions with Dlx transcription factors, a mechanism previously described for Sp7 in bone development (Hojo et al., 2016). Notably, several Dlx family members, including Dlx5 and Dlx6, share expression domains with Sp6 and Sp8 in the limb ectoderm. Furthermore, the *Dlx5/Dlx6* double knockout embryos present SHFM phenotype, resembling that of *Sp6;Sp8* compound mutants retaining a functional *Sp8* allele (Haro et al., 2014; Robledo et al., 2002).

To investigate whether Sp8 and Sp6 physically interact with Dlx5, we performed co-immunoprecipitation (CoIP) assays in HEK293 cells co-expressing epitope-tagged (FLAG or Myc) of these factors. These assays confirmed that Dlx5 interacts with both Sp8 and Sp6 and that this interaction requires the ZF domain of Sp proteins (Fig. 5I), similar to what was previously reported for Sp7 (Hojo et al., 2016).

To corroborate the cooperative binding of Sp6 and Sp8 to AT-rich motifs via interaction with Dlx5, we performed an additional EMSA using the probe containing the AT-rich/Dlx5-binding motif. As expected, Dlx5 (Myc-tagged) alone formed a shifted complex, which was supershifted upon addition of an anti-Myc antibody, confirming the identity of the complex (Fig. 5J, lanes 2–3). Co-expression of Sp6 (FLAG-tagged) or Sp8 (FLAG-tagged) with Dlx5 resulted in a distinct shifted band, which was further supershifted (in the case of Sp6) or no longer detected (in the case of Sp8) upon addition of the anti-FLAG antibody. These changes indicate that both Sp6 and Sp8 are incorporated into the Dlx5-DNA complex (Fig. 5J, lanes 4–7). These findings support a model in which Sp6 and Sp8 do not directly bind AT-rich motifs but are recruited to these regulatory elements through protein–protein interactions with Dlx cofactors. Protein expression levels of Sp6, Sp8, and Dlx5 were quantified by Western blot analysis (Fig. S4D).

The fact that Sp8 utilizes the same domain, the ZF, to recognize its cognate DNA sequence and to mediate protein interactions highlights the complexity and context-dependence of its regulatory function. Collectively, our results support a model in which Sp8, Sp6 and Dlx5 collaboratively regulate target genes with the final functional outcome depending on the relative abundance of each transcription factor.

## DISCUSSION

Here, we uncover the complex regulatory networks by which Sp8 and Sp6, cooperatively and independently, integrate Wnt/β-catenin, FGF, and BMP signaling, to fine-tune gene expression during limb development.

To determine the molecular mechanisms underlying Sp8 and Sp6 transcriptional activity in the limb ectoderm at a genome-wide scale, we used ChIPmentation and RNA-seq. To distinguish between cooperative and specific regulatory roles, we overlapped the chromatin binding profiles of Sp8 and Sp6 and classified the binding regions in three categories: peaks co-occupied by both factors, and peaks specifically bound by either Sp8 or Sp6. To distinguish functional from non-functional binding events, we intersected these peak categories with the sets of DEGs identified upon the individual deletion of Sp8 or Sp6. Genes associated with regulatory regions co-occupied by both transcription factors and showing altered expression in either one or both single mutants were considered as potential common targets. Among these, 12 genes required both factors for proper expression, considered genuine common targets, while others depended solely on Sp8 (119) or on Sp6 (20) for normal expression. We also identified direct target genes specific to Sp8 and Sp6, each associated with regions bound exclusively by one of these factors.

Our data also reflect the dominant transcriptional role of Sp8 compared to Sp6, consistent with the more severe phenotype of *Sp8* mutants. One contributing factor to this difference may be that *Sp6* is a Sp8 direct target. We previously showed that *Sp6* expression is not maintained in *Sp8* mutants (Haro et al., 2014) and here we identify a putative regulatory element within the *Sp6* gene through which Sp8 exerts this control. As *Sp6* expression declines, *Sp8* mutants may effectively function as double mutants. Additionally, Sp8’s unique ability to bind GC-rich proximal regions, where Sp6 cannot compensate, further explains its dominance.

### The Complexity of Sp8/Sp6 Regulatory Network in the Limb Bud Ectoderm

Sp8 is a well-recognized mediator of the Wnt/βcatenin dependent induction of *Fgf8* in the limb ectoderm and also in other systems such as the genital tubercle and the embryonic telencephalon (Haro et al., 2014; Lin et al., 2013; Sahara et al., 2007). Together with Sp5, Sp8 is also a mediator of Wnt/βcatenin dependent induction of neuromesodermal progenitors (Kennedy et al., 2016). However, we did not detect significant enrichment of Tcf/Lef motif in our ChIPmentation data, suggesting that in the limb ectoderm Sp8/Sp6 do not operate by binding to βcatenin and rather mediate Wnt/β catenin signaling by regulating the expression of key pathway components such as *Sost*, *Rspo2* and *Fzd1*, which are regulated by common distal regulatory elements and dependent on both (*Sost*) or solely on Sp8 (*Rspo2* and *Fzd1)*. Thus, by coordinating both positive and negative components of the Wnt/β-catenin pathway Sp6/Sp8 ensure proper signaling dynamics during limb development. Similarly, our ChIPmentation data showed no enrichment of Smad motifs, suggesting Sp6/Sp8 modulate Bmp signaling by regulating *Bmp4*, as well as its bona fide effectors *Msx1* and *Msx2*.

In addition to modulating key signaling pathways, Sp8 and Sp6 also directly regulate crucial patterning genes in the limb ectoderm including *Fgf8* and the DV core factors *En1* and *Wnt7a*. Here, we show that Sp8, but not Sp6, directly regulates *Fgf8*, while both Sp8 and Sp6 contribute to the regulation of *En1* and *Wnt7a* with higher contribution of Sp8.

*Fgf8* expression, the marker of the AER, is differentially affected in *Sp* mutants. It is initiated but not maintained in *Sp8* mutants, remains unaffected in the double ridge of *Sp6* mutants, and is never activated in *Sp8;Sp6* double mutants. Here, we show that Sp8 directly regulates *Fgf8* by binding to GC-rich motifs within its promoter. In contrast, we did not identify any *Sp6*-bound regulatory regions associated with *Fgf8*, suggesting that the Sp6 input observed in double mutants is indirect, potentially mediated through its role in modulating Wnt signaling in the ectoderm. In addition to promoter regulation, *Fgf8* expression is governed by multiple enhancers located in a large regulatory region centromeric to the gene (Hörnblad et al., 2021; Marinić et al., 2013). Duplications of this genomic region have been linked to the SHFM3 syndrome (OMIM 246560), which is thought to result from disrupted genomic architecture and altered AER expression (Cova et al., 2023). However, our analysis found no evidence of Sp8 or Sp6 binding to this complex regulatory domain.

We also show that *En1* and *Wnt7a* possess distal enhancer co-bound by Sp8 and Sp6, but requiring only Sp8 for normal expression (Fig. 4A). The *En1* enhancer which we show drives activity in the apical ectodermal ridge (AER) in mouse transgenic assays and overlaps the first exon of the *En1*-associated long non-coding RNA (lncRNA) *Lmer* (Ringel et al., 2024). However, deletion of the *Lmer* locus, including the Sp8/Sp6-bound enhancer, did not affect *En1* expression in the limb ectoderm. Thus, despite its ability to drive expression to the AER and ectoderm, this enhancer is dispensable for *En1* function, likely due to redundancy with other elements such as Maenli. This suggests that the downregulation of *En1* observed in *Sp8* and double mutants must depend on additional factors, possibly Bmp4, which could also explain the delayed onset of this effect.

Our study also identifies a Sp8/Sp6 co-bound enhancer that positively regulates *Wnt7a*. This may appear unexpected, as *Wnt7a* expression extends ventrally in both *Sp8* and *Sp8;Sp6* mutants. However, this expansion is indirect, in parallel to the loss of *En1*. When expression levels are considered, both ISH and RNA-seq reveal that *Wnt7a* is reduced in *Sp8* and *Sp8;Sp6* mutants strongly suggesting that its expression is directly regulated thought the Sp8/Sp6 bound enhancer. A major question that remains open is how Sp8 and Sp6, initially expressed throughout the entire limb ectoderm, contributes to the establishment of the defined DV expression domains.

These findings highlight Sp8 and Sp6’s role in coordinating PD and DV patterning and underscore the complexity of their cooperative and independent function.

### Sp8 Has a Dual Mode of Action

Another key finding of our study is that Sp8 and Sp6 operate through distinct mechanism. Motif analysis, EMSA and co-IP experiments, revealed that Sp8 uses a dual strategy: it binds DNA directly at GC-rich motifs and indirectly at AT-rich regions, likely via interactions with Dlx proteins. In contrast, Sp6 appears to rely primarily on indirect DNA binding via co-factors, predominantly Dlx proteins. These differences challenge the previously proposed functional redundancy between Sp8 and Sp6 suggested by earlier genetic studies in mice (Haro et al., 2014). Furthermore, our results show that Sp8 preferentially binds GC-rich motifs when acting independently, whereas its cooperation with Sp6 is associated with binding at AT-rich sites, a subject that merits further investigation. Although we demonstrate that Sp8 and Sp6 can form heterodimers, this is likely not the dominant in vivo scenario, given the limited overlap in their binding sites, possibly due to interactions with distinct co-factors.

Another member of the Sp family, Sp7, is unable to bind the canonical Sp consensus motif and instead engages DNA through interaction with Dlx5 (Hojo et al., 2016). This distinct mode of action is attributed to three specific amino acid substitutions in its ZF domain, which impair GC-box recognition while enhancing Dlx binding. However, Sp8 and Sp6 do not share these Sp7-specific variants, indicating that these residues are not required for Dlx5 interaction. Interestingly, Sp9, another Sp protein expressed in the ectoderm, shows a similar binding pattern to that of Sp8, as published motif enrichment analysis in ganglionic eminences indicate: AT-rich (Xu et al., 2018) and GC-box (Catta-Preta et al., 2025). Our findings support the idea that interaction with Dlx is a more broadly conserved feature within the Sp family, at least among members of the Sp6–Sp9 clade (Fig. 6).

**Figure 6.**
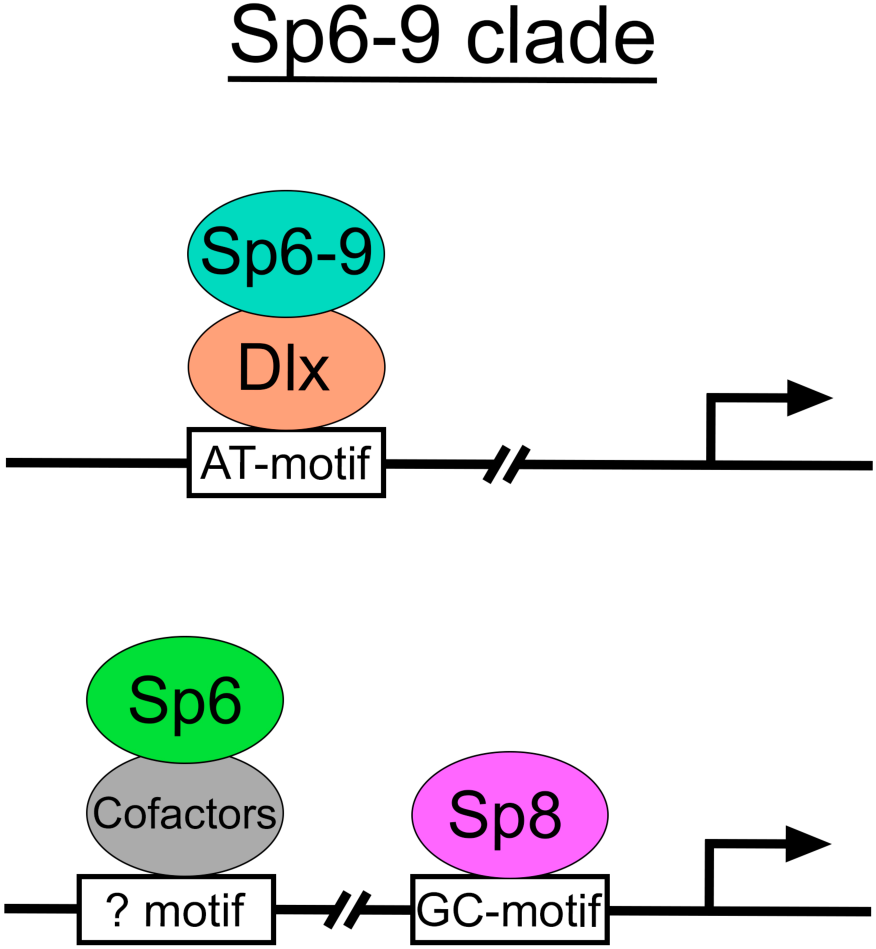
Working model illustrating the mode of action of the Sp6-Sp9 clade in the limb ectoderm.

### Impact in Human Congenital Malformation

Progressive disruption of the Sp8/Sp6 gene regulatory network in mice results in a spectrum of limb malformations, including oligodactyly, ectrodactyly, and amelia, mirroring human congenital disorders. In particular, the *Sp8+/-;Sp6-/-* mutant serves as a robust model for human SHFM (Haro et al., 2014). Although no pathogenic variants in human *SP8* or *SP6* have been identified in SHFM cases, their mechanistic interaction with DLX5 suggest a potential implication in SHFM type 1, associated to *DLX5/6* and in SHFM type 5, associated to *DLX1/2*. Furthermore, *Bhlha9*, a common transcriptional target of Sp8 and Sp6, is linked to SHFM with long bone deficiency 3 (SHFLD3) suggesting a potential role for Sp8/Sp6 in thie pathology.

The complexity of the Sp8/Sp6 network may rely on their ability to interact with each other as well as with other TFs. Cofactor availability may influence their mechanistic mode of action. In the limb ectoderm, Sp8, Sp6, and Dlx5—along with other members of their respective families—likely form dynamic regulatory assemblies whose composition and activity depend on context, affinity, and expression levels. Adding another layer of complexity, Sp8 directly activates the transcription of several *Dlx* genes and *Sp6*, establishing autoregulatory and feedforward circuits.

In conclusion, our study identifies Sp8 and Sp6 as key regulators of gene expression in the limb ectoderm, functioning both independently and cooperatively. Collectively, our results support a model in which Sp8, Sp6, and Dlx5 collaboratively regulate target genes, with the final functional outcome depending on the relative abundance of each transcription factor. These findings highlight the central role of Dlx– Sp interactions in orchestrating limb morphogenesis and underscore the importance of considering transcription factor networks, rather than individual factors, in the study of congenital malformations.

## MATERIAL AND METHODS

### Mouse Embryos

Wild type C57BL/6 mice and the *Sp6^-/-^* (Nakamura & Ishikawa, 2008), *Sp8^FL^* and S*p8^CreERT2^* (Treichel et al., 2003) mouse strains were used in this study. Genotyping was performed using tail biopsies or embryonic membranes according to previously published reports (Haro et al., 2014). Embryos of the desired embryonic day were obtained by cesarean section. All animal procedures were conducted accordingly to the EU regulations and 3R principles and reviewed and approved by the Bioethics Committee of the University of Cantabria. All mice were maintained in a C57BL6 genetic background and genotyped by PCR protocols from tail biopsies or yolk sacs samples following standard protocols.

### *Sp8:3xFLAG* Knock-in Mouse Generation (*Sp8^FL^*)

Three copies of the FLAG epitope (5’-GACTACAAAGACCATGACGGTGATTATAAAGATCATGATATCGATTACAAGGATGACGATGACAAG-3’) were inserted in frame at the 3’-terminal end of the *Sp8* gene with a stop codon (Fig. 1A). Mouse genomic fragments containing 5’ (chr12:118845063+118849869) and 3’ (chr12:118849873+118852808) homology arms were amplified from BAC clone by using high fidelity Taq polymerase and were sequentially assembled into a targeting vector together with recombination sites and selection markers (DTA and Neo cassette). Targeted ES clones were confirmed via Southern Blotting and some of them were selected for blastocyst microinjection (Cyagen Biosciences Inc., Santa Clara, California, US).

### Isolation of Limb Bud Ectodermal Hulls

E10.5 forelimb buds were dissected in cold PBS and incubated in 0.25% trypsin (SV30037.01, GE Healthcare, Logan, Utah) on ice for 20 min. After a quick rinse in 10% FBS to inactivate trypsin, the limb buds were transferred to cold PBS where the ectoderm was separated from the mesoderm with fine forceps.

### Transgenic Mice for LacZ activity

The genomic region containing the regulatory elements of *En1* (mm10, chromosome 1:120,621,403-120,622,419) and *Rspo2* (mm10, chromosome 15:43,388,035-43,389,518) were PCR-amplified using specific primers containing XhoI restriction enzyme sites and cloned into a vector carrying the β-globin minimal promoter and the LacZ coding sequence (pSK-LacZ; kindly provided by Denis Duboule lab). Transient transgenic embryos were harvested at E10.5 and only those embryos containing the reporter transgene were processed for detection of β-galactosidase activity according to standard procedures.

### Skeletal Preparations, *in situ* Hybridization (ISH) in Paraffin Sections and Immunofluorescence

Alcian blue 8GX and Alizarin red skeletal staining was performed following standard protocols, cleared by KOH treatment and stored in glycerol. *In situ* hybridization was performed on paraffin sections with digoxigenin-labeled antisense riboprobes following standard procedures. Immunostaining was performed in cryostat sections (14 µm) using the mouse monoclonal anti FLAG M2 antibody (1:500; Ref F1804; Sigma-Aldrich, St Louis, Missouri) and the anti Sp8 C18 antibody (1:500; Ref 104661, Santa Cruz Biotechnology, Dallas, Texas) and then coupled with Alexa 488-conjugated anti-mouse IgG secondary antibody (1:500, Invitrogen, Molecular Probes, Eugene, Oregon). Slides were analyzed in a Leica Laser Scanning Confocal TCS-SP5 with a 63x 1.4 NA objective.

### Plasmids Constructions

Sp8 and Sp6 tagged with FLAG or Myc epitopes at their 5’ end were amplified by PCR and cloned into pcDNA3 including the appropriate restriction enzyme site to be fused at the 3’end to the full length YFP (1-240aa) or to its N-terminal (YN; 1-172aa) or C-terminal (YC; 173-240aa) moieties. Deletion of the ZF domain of Sp8 (nt 353-486) and of the ZF domain of Sp6 (nt 254-336) was constructed by PCR strategy. The following clones were generated: FLAG-Sp8, FLAG-Sp6, HA-Sp6ΔZF, Sp8-YFP, Sp8-YN, Sp8-YC, Myc-Sp6-YFP, Myc-Sp6-YN, Myc-Sp6-YC, Myc-Sp8ΔZF-YFP, Myc-Sp8ΔZF-YN, Myc-Sp8ΔZF-YC. The Myc-Dlx5 and FLAG-Dlx5 clones were kindly provided by Dr. Andrew P. McMahon (Keck School of Medicine of the University of Southern California, Los Angeles, USA) (Hojo et al., 2016). The HA-Sp6ΔZF clone was synthesized by NZYTech (Lisbon, Portugal).

### Bimolecular Fluorescent Complementation (BiFC)

HEK293 cells were grown on microscope cover Glasses (Ø 18 mm) and transfected with PEI and 1 µg total DNA. To test for homodimerization, 500ng of indicated expression plasmids, each one with a complementary YFP moiety (e.g. Sp8-YN + Sp8-YC), were transfected. Single transfection of the protein fused with the complete YFP or with one of its moieties was used for positive and negative control. Transfected, cells were incubated with Hoechst33342 and confocal images (512×512 pixels; 0.15mm pixel size) were acquired sequentially on a SP5 laser-scan microscope (Leica) with a 63x 1.4 NA objective. Cells were excited sequentially with 405nm 514nm laser lines and fluorescence emission captured between 420-480nm (Hoechst) and 525-600nm (YFP). Images are presented after digital adjustment of curve levels (gamma) to maximize signal with ImageJ software. Fluorochromes and colors are as indicated in the figure legends. Images are representative of at least three independent experiments.

HEK293 cells were grown on microscope cover Glasses (Ø 18 mm) and transfected with PEI and 1 µg total DNA. To test for homodimerization, 500ng of indicated expression plasmids, each one with a complementary YFP moiety (e.g. Sp8-YN + Sp8-YC), were transfected. Single transfection of the protein fused with the complete YFP or with one of its moieties was used for positive and negative control. Transfected, cells were incubated with Hoechst33342 and confocal images (512×512 pixels; 0.15mm pixel size) were acquired sequentially on a SP5 laser-scan microscope (Leica) with a 63x 1.4 NA objective. Cells were excited sequentially with 405nm 514nm laser lines and fluorescence emission captured between 420-480nm (Hoechst) and 525-600nm (YFP). Images are presented after digital adjustment of curve levels (gamma) to maximize signal with ImageJ software. Fluorochromes and colors are as indicated in the figure legends.

### Co-immunoprecipitation, SDS-PAGE, and Western Blotting

HEK293 cells were co-transfected with the desired plasmids (1 µg total DNA) and lysed (20 mM Hepes (pH 7.5), 10 mM EGTA pH 8, 40 mM Glycerol-phosphate, 1% NP-40, 2.5 mM MgCl2, 2 mM orthovanadate, 1 mM DTT, 1x PICS, 1 mM PMSF). CoIP was performed with 0.5-1mg of protein, 10 µL of Dynabeads™ Protein G (Thermo Fisher Scientific, 10004D) and 1μg of the desired antibody: FLAG-M2 (Sigma F1804) or MYC (Cell Signaling #2272) followed by SDS-PAGE analysis of protein products. FLAG-M2 and MYC antibodies were used for immunoblotting and β-Actin (Santa Cruz #47778) or H3 (Cell Signaling #96C10) was used as loading control when needed. Results were confirmed in at least two independent experiments.

HEK293 cells were co-transfected with the desired plasmids (1 µg total DNA) and lysed (20 mM Hepes (pH 7.5), 10 mM EGTA pH 8, 40 mM Glycerol-phosphate, 1% NP-40, 2.5 mM MgCl2, 2 mM orthovanadate, 1 mM DTT, 1x PICS, 1 mM PMSF). CoIP was performed with 0.5-1mg of protein, 10 µL of Dynabeads™ Protein G (Thermo Fisher Scientific, 10004D) and 1μg of the desired antibody: FLAG-M2 (Sigma F1804) or MYC (Cell Signaling #2272) followed by SDS-PAGE analysis of protein products. FLAG-M2 and MYC antibodies were used for immunoblotting and β-Actin (Santa Cruz Ref 47778) was used as loading control when needed. Results were confirmed in at least two independent experiments.

### EMSA Assay

EMSA was performed using the nuclear extracts from HEK-293 cells or HEK-293 cells transfected with Sp8, Sp6 and Dlx5. DNA probes labeled with IRDye700 on 5’end were synthesized by Integrated DNA Technologies, Inc. (IDT, Coralville, IA). The sequences of probes are shown in (Hojo et al., 2016). Each reaction was performed by mixing 15 ug of the nuclear extract with the binding buffer (250 mM KCl, 100 mM HEPES-KOH (pH 7.5), 0.875 mM EDTA, 25% glycerol, 1.5 μg of Poly d(I-C)). Mixtures were pre-incubated at RT for 15 min. Later 0.25 mM of the Biotin-labeled probe was added, and samples were incubated 15 min at RT. For supershift assay, 1 ug of anti-FLAG M2 antibody (Sigma-Aldrich, F1804) or anti-MYC (Cell Signaling #2272) was incubated at RT for 15 min prior to binding with probes. For EMSA involving the Sp–Dlx complex (Fig. 5K), all the incubations steps were extended to 30 minutes. Complexes were analyzed by electrophoresis on a 6% polyacrylamide gel in 0.5x TBE buffer and visualized using an Odyssey infrared imaging system (Li-Cor Bioscience). Results were validated in at least two independent experiments.

EMSA was performed using the nuclear extracts from HEK-293 cells or HEK-293 cells transfected with SP8 and SP6. DNA probes labeled with IRDye700 on 5’end were synthesized by Integrated DNA Technologies, Inc. (IDT, Coralville, IA). The sequences of probes are shown in (Hojo et al., 2016). The reaction was performed by mixing 15 ug of the nuclear extract with the binding buffer (250 mM KCl, 100 mM HEPES-KOH (pH 7.5), 0.875 mM EDTA, 25% glycerol, 1.5 μg of Poly d(I-C)). Mixtures were pre-incubated at RT for 15 min. Later 0.25 mM of the Biotin-labeled probe was added to each condition and the resulting mixture was incubated 15 min at RT. For supershift assay, 1 ug of anti-FLAG M2 antibody (Sigma-Aldrich, F1804) was incubated at RT for 15 min prior to binding with probes. Complexes were analyzed by electrophoresis on a 6% polyacrylamide gel in Tris-Borate-EDTA buffer 0.5x. Geles were visualized using an Odyssey infrared-imaging system (Li-Cor Bioscience).

### Luciferase Reporter Assay

NIH 3T3 cells were plated at a density of 25,000 cells/cm2 (22Rv1) in 24-well plates and the next day were transfected by Lipofectamine^TM^ 3000 (L3000-008 Invitrogen by Thermo Fisher Scientific Life Technologies, Carlsbad, California) with 50 ng/well of firefly luciferase reporter SP1 sites (GC-box) (93p21-luc wt or 93p21-luc(mut 2+3+4) (Vaqué et al., 2005), 15 ng of the pRL-TK Renilla luciferase control plasmid (Promega) and 50ng of FLAG-Sp8 or FLAG-Sp6 expression plasmids. For dose-response experiments, 12.5 ng and 25 ng of FLAG-Sp6 plasmid were also tested. After 48 hours, luciferase activity was measured with the DualGlo Stop&Glo Luciferase Assay System ( Promega #E2920), following the manufacturer’s instructions. Each experiment was performed in duplicate and was repeated at least three times. Error bars represent standard deviations calculated from the biological triplicates.

Expression profile of NIH 3T3 cells were obtained from BioGPS (http://amp.pharm.mssm.edu/Harmonizome/gene_set/nih+3T3/BioGPS+Mouse+Cell+Type+and+Tissue+Gene+Expression+Profiles).

### ChIPmentation

ChIPmentation protocol was based on (Schmidl et al., 2015). In order to reduce background and unspecific binding, only the limb bud ectoderm was used. A total of 50 ectoderms from *Sp8*^FLAG/FLAG^ (for Sp8) and WT (for Sp6) were used for each TF ChIPmentation replicate (2 replicates). ChIPmentation sequencing data was processed by a standard workflow using nf-core/chipseq/2.1.0 pipeline (Ewels et al., 2020). Because of the higher signal to noise ratio, only the dataset of the first biological ChIPmentation sample was used for further analysis. For proximal and distal classification, distance to the nearest tss was obtained using bedtools closest function (Quinlan & Hall, 2010). Pearson Correlation coefficients of the two SP8 ChIPmentation biological replicates were performed with the bamCorrelate tool from deepTools (bins mode and a bin size of 10 kb across the whole mouse genome) (Ramírez et al., 2014). For the generation of the heatmaps, BAM files were normalized as RPGC (reads per genome coverage) and then used to visualize scores associated with genomic regions using deepTools (Ramírez et al., 2014). The bigwig of scATAC-seq ectoderm data was obtained from the GEO accession number GSE: (Desanlis et al., 2020). Average vertebrate PhastCons score profiles around the center of enhancer sequences were generated with the Conservation Plot tool from the Cistrome Analysis pipeline (http://cistrome.dfci.harvard.edu/ap/root) (Liu et al., 2011).

For the clustering analysis, Sp8 and Sp6 peaks were concatenated and merged with bedtools merge function. Then, k-means ranked clustering of peaks based on Sp8 and Sp6 ChIPmentation signal was performed with seqMINER 1.3.4 (Ye et al., 2011), targeting 12 clusters and with the seed 13,625,001. Clusters with similar profile were merged to obtain 3 final clusters: Common, Sp8-, and Sp6-predominant.

### RNA-seq

Ectoderms from E10.5 wild-type and homozygous *Sp8-*null (*Sp8^Cre-ERT2^* line) and Sp6-null (Sp6KO line) embryos were individually collected. Experiments were performed in triplicates and each replicate contained the 2 forelimb ectoderms of a single embryo. RNA was extracted with RNeasy-Plus Micro Kit (Ref 74034 Qiagen GmbH, Hilden, GERMANY). Pre-amplification using the Ovation RNASeq System V2 was performed. Total RNA was used for first strand cDNA synthesis, using both poly(T) and random primers, followed by second strand synthesis and isothermal strand-displacement amplification. For library preparation, the Illumina Nextera XT DNA sample preparation protocol was used, with 1 ng cDNA input. After validation (Agilent 2200 TapeStation) and quantification (Invitrogen Qubit System) all six transcriptome libraries were pooled. The pool was quantified using the Peqlab KAPA Library Quantification Kit and the Applied Biosystems 7900HT Sequence Detection and pooled on one lane of an Illumina HiSeq4000 sequencing instrument with a 2×75 bp paired-end read length.

To analyze the data, high-throughput next-generation sequencing analysis pipeline (Wagle et al., 2015) was used. Basic read quality check was performed using FastQC (Babraham Bioinformatics) and reads filtered and adaptor-trimmed with fastp (Chen et al., 2018). Reads were mapped to the mouse reference assembly (GRCm38), using HiSat2 (Kim et al., 2019) with the option --dta-cufflinks, which enabled direct transcript assembly with StringTie (Pertea et al., 2015). Read count means, fold-change (FC) and values were calculated with DEseq2 (Anders & Huber, 2010). Genes were considered significant differentially expressed genes when p-adjusted value < 0.05 and log2FC > 0.585 (FC > 1.5).

Sp6 expression was not properly calculated by the computational pipeline described above due to the presence of an antisense transcript that largely overlaps with Sp6 and that got assigned most of the RNA-seq reads (i.e. ENSMUSG00000087067). Since according to the RNA-seq data ENSMUSG00000087067 was downregulated in *Sp8KO* embryos and our RNA-seq data was not strand-specific, this indicates that Sp6 is indeed downregulated in *Sp8KO* embryos. This was further confirmed by ISH and is also supported by our previous findings (Haro et al., 2014). Therefore, the read values measured for ENSMUSG00000087067 were assigned to *Sp6* instead.

Gene ontology was performed with the DAVID web-based tool (davidbioinformatics.nih.gov, (Sherman et al., 2022)) with default parameters.

### Motif Analysis

*De novo* motif discovery analyses of Sp8 peaks were performed using the online tool MEME-ChIP (Version 5.0.1, http://meme-suite.org/tools/meme-chip) as described by (Ma et al., 2014). Input sequences were centered within summit regions of recovered intervals obtaining motifs that are between 6 and 20 bp wide with an E-value cut-off of >0.5 for the discovery of enriched motifs.

The database used to identify known motifs was HOCOMOCO H12 CORE (human + mouse) (Kulakovskiy et al., 2018). For the prediction of individual motif occurrences, FIMO tool from the meme suite with default parameters. The AT-rich and GC-box motifs obtained from MEME-ChIP with Sp8 peaks were used as input.

### Genomic Regions Enrichment of Annotations Tool (GREAT) analysis

For calculating the distribution of all the Sp8 peaks with respect to the TSS of annotated genes as well as for their *in silico* functional annotation analyses, we used the GREAT tool (GREAT 3.0.0) (McLean et al., 2010) with the “Basal Plus Extension Association Rule” settings. *Rspo2* was assigned to the peak chr15:43388486-43388897 given their proximity, and was subsequently validated with Lacz transgenic reporter analysis.

## Acknowledgments

We are very grateful to Dr. Hojo (University of Tokyo) and Dr. McMahon (University of Southern California) for the p12xAT_luc, the p12xAT_luc and Dlx5 constructs (Hojo et al., 2016), to Susan Mackem and Juan Tena for discussions. Work in the Rada-Iglesias laboratory is supported by grants PID2021-123030NB-I00, ERC CoG PoisedLogic (862022) and Chrom_rare HORIZON-MSCA-2021-DN-01-101073334. AC-I and SM were recipients of PhD fellowships PRE2018-083421, PRE2021-098480 respectively from the Spanish Ministry of Science and Innovation. This research was supported by the Spanish Ministry of Science and Innovation PID2023-147771NB-I00 to MAR.

## Author contributions

A.C-I, S.M. and R.P-G performed the experiments. V.C. contributed to experiments. A.R-I. and J.F.L-G. contributed to the design of the project and the analysis of the data. M.A.R. conceived the project, performed experiments, analyzed the data and wrote the manuscript. All the authors edited the manuscript.

## Data and materials availability

The original ChIPmentation and RNA-seq data of this paper have been deposited in GEO and can be accessed with the accession number XXX (available to reviewers upon request and will be made public upon publication).

## Source Data 1-5

Source Data 1-Sp8 and Sp6 ChIPmentation peaks (Excel)

Source Data 2-In silico functional annotation of Sp8 and Sp6 peaks using GREAT (Excel)

Source Data 3-Differentially expressed genes between WT and *Sp8*- and *Sp6-* null embryos and corresponding Gene Ontology analysis (Excel)

Source Data 4-Peaks classified by category and intersections with DEGs (Excel)

Source Data 5-Gene Ontology analysis of different categories (Excel)

