## Supplementary Figures for "*Sp8* and *Sp6* regulatory functions in the limb bud ectoderm"

### SUPPLEMENTARY FIGURE 1

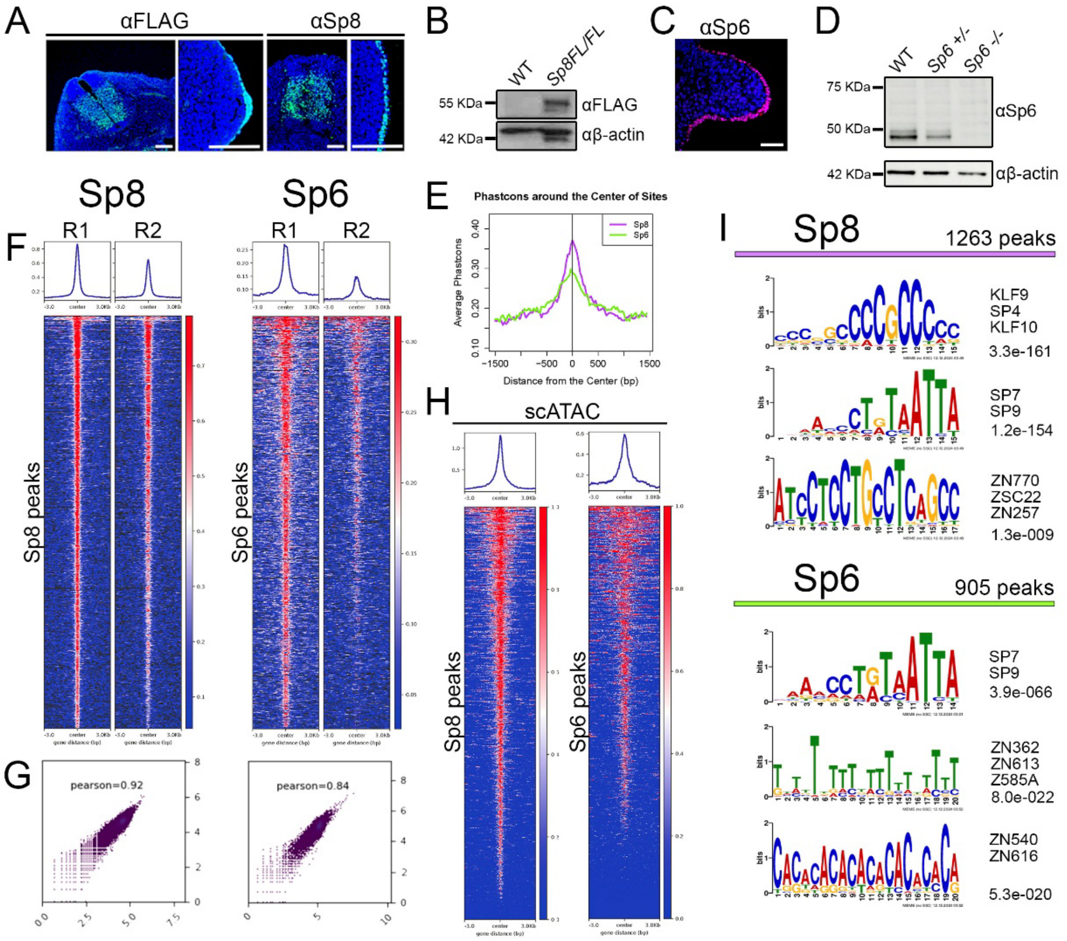


**Figure S1. Validation of Sp8-3xFLAG knock-in mice, Sp6 antibody and of the Sp8/6 ChIP-seq data.**

(**A**) Immunofluorescence with anti-FLAG, and with anti Sp8 antibodies in cryostat sections of an E10.5 Sp8FL/FL embryo showing similar expression pattern in the neural tube and in the limb ectoderm. **(B**) Western blot showing detection of Sp8FL in Sp8FL/FL but not in WT E10.5 embryos. β- actin loading control is shown below. **(C**) Immunofluorescence anti-Sp6, in cryostat sections of an E10.5 WT embryo. **(D)** Western blot validating the specificity of the anti-Sp6 antibody in E10.5 limb buds. β-actin, as a loading control, is shown below. **(E**) Average vertebrate PhastCons score profiles around the center of Sp8 (purple line) and Sp6 (green line) binding regions showing evolutionarily conservation. **(F**) Average profile and heatmaps of Sp8 and Sp6 ChIPmentation peaks in the two biological replicates (R1 and R2). Represented regions are +3 kb and -3 kb from the peak center. **(G)** Pearson correlation between the two ChIPmentation replicates for Sp8 and Sp6. **(H)** Average profile and heatmaps of the identified Sp8 and Sp6 peaks in ectoderm ATAC-seq data obtained from an available limb scATAC-seq dataset. (**I**) Top three most significant motifs enriched at Sp8 and Sp6 ChIPmentation peaks identified by *de novo* motif analysis with MEME-ChIP. Best matches with known motifs and the motif e-value are shown at the right side.

#### SUPPLEMENTARY FIGURE 2

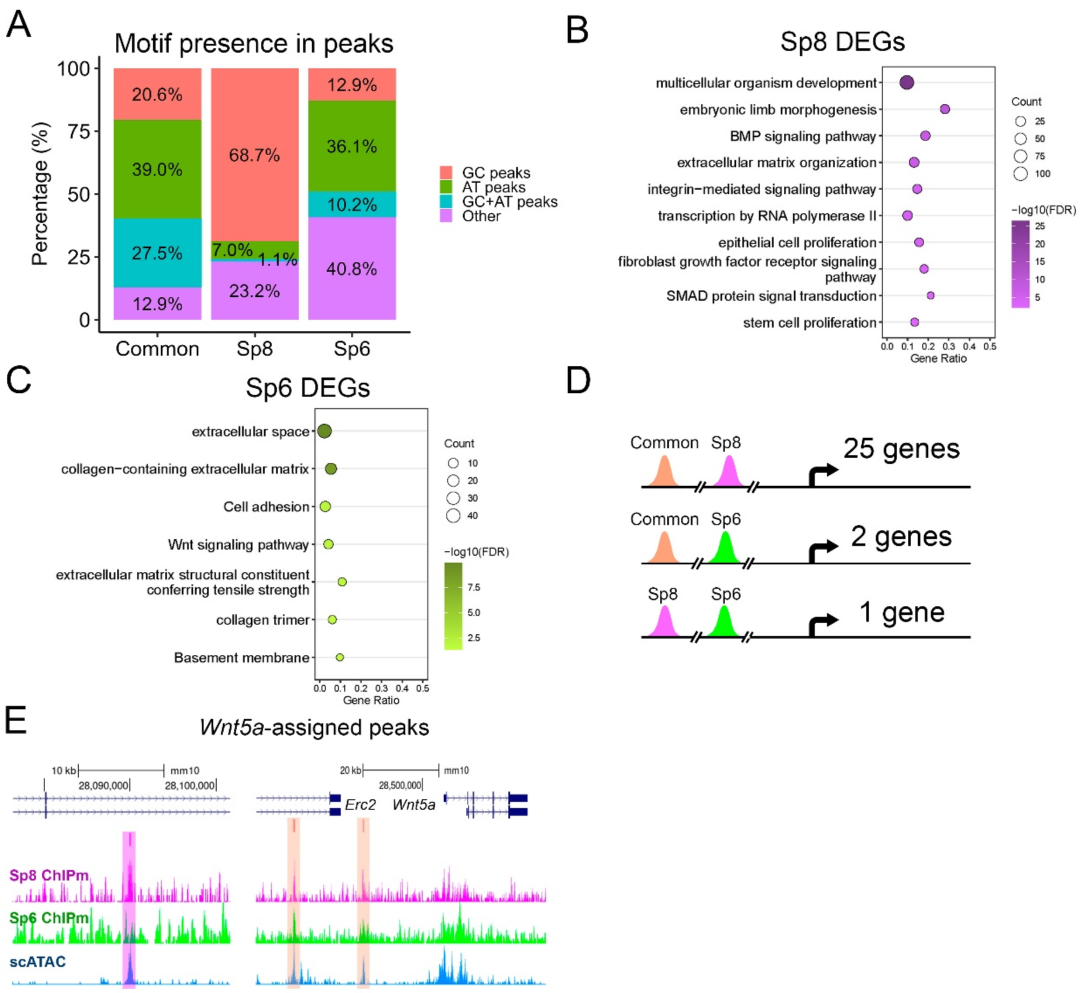


**Figure S2. Characterization of Sp8 and Sp6 target regions and gene regulatory programs.** **(A)** Distribution of motif types within peaks assigned to common, Sp8-specific, or Sp6-specific peaks. Peaks were categorized as containing only GC-rich, only AT-rich, both GC+AT, or containing other motifs. **(B-C)** Gene Ontology (GO) enrichment analysis of differentially expressed genes (DEGs) in Sp8 (B) and Sp6 (C). **(D)** Schematic showing the number of genes with regulatory regions bound by common, Sp8-specific, or Sp6-specific peaks. A total of 25 genes are associated with common and Sp8 peaks, 2 with common and Sp6 peaks, and 1 gene (*Lgr6*) with both Sp8- and Sp6-specific peaks. **(E)** Genome browser tracks showing peak assigned to *Wnt5a*. Tracks display peaks for common (orange), Sp8-specific (pink), and Sp6-specific (green) peaks.

#### SUPPLEMENTARY FIGURE 3

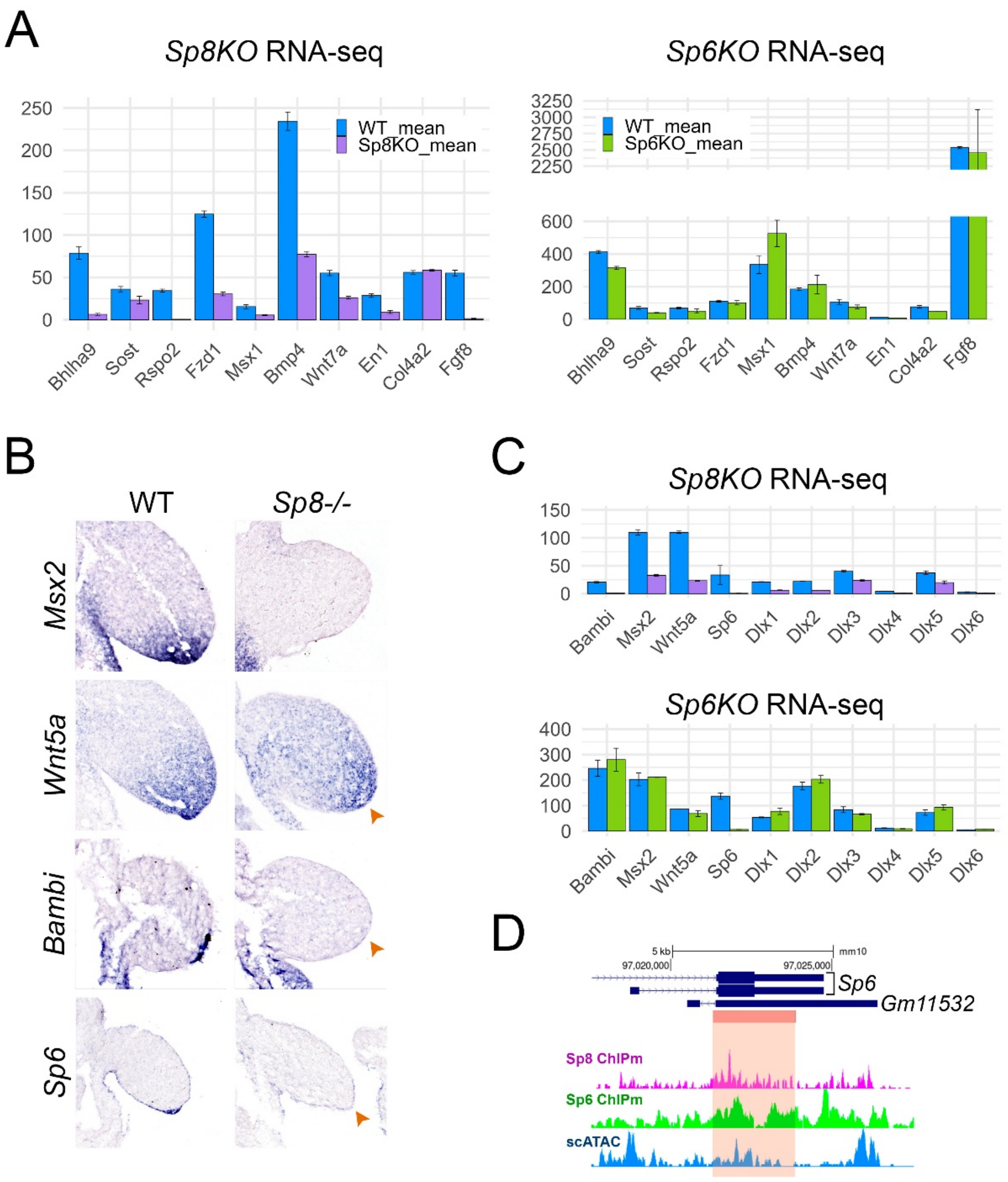


**Figure S3. Validation of Sp8/6 targets**
**(A)** Bar graphs showing RNA-seq expression levels (mean ± SD) of candidate target genes in *Sp8-/-* (left) and *Sp6-/-* (right) limb buds compared to WT controls. **(B)** ISH of *Msx2*, *Wnt5a*, *Bambi*, and *Sp6* in WT and *Sp8-/-* E10.5 limb buds reveals reduced expression in mutants (orange arrowhead), consistent with RNA-seq results. **(C)** Additional RNA-seq expression bar graphs of target genes in *Sp8-/-* (top) and *Sp6-/-* (bottom) limb buds compared to WT controls. **(D)** Screenshot from UCSC genome browser showing the common intragenic peak associated to Sp6 gene. *Gm11532*, the antisense transcript to whom all the Sp6 reads were assigned is also shown.

#### SUPPLEMENTARY FIGURE 4

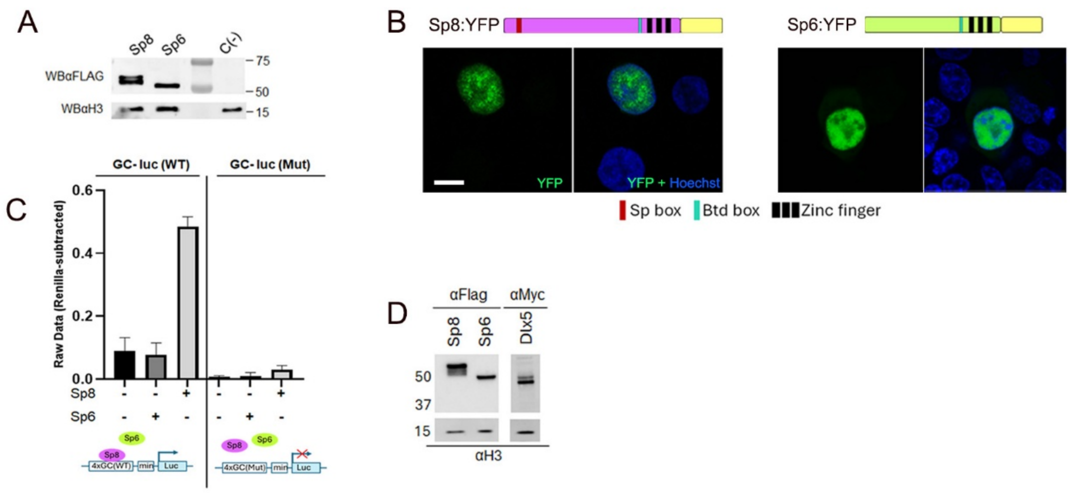


**Figure S4. Functional analysis of Dlx5, Sp8 and Sp6 interactions and nuclear localization of Sp8 and Sp6.**

**(A)** Western blot analysis of the nuclear extracts used in Figure 5A–B, confirming comparable protein expression levels across samples. **(B)** BiFC assay shows nuclear localization of Sp8 and Sp6 in HEK293 cells. Transfected construct indicated at the top. Scale bar: 10 µm. **(C)** Luciferase reporter assay using constructs containing WT or mutant GC-box motifs. Sp8 activates the WT GC-box reporter, while Sp6 alone has no effect. Mutation of the GC-box abolishes Sp8-mediated activation. Error bars represent standard deviations calculated from the biological triplicates. **(D)** Western blot analysis of the nuclear extracts used in Figure 5J, confirming comparable protein expression levels.
